# Different protein metabolic strategies for growth during food-induced physiological plasticity

**DOI:** 10.1101/2020.06.07.139139

**Authors:** Aimee Ellison, Amara Pouv, Douglas A. Pace

## Abstract

Food-induced morphological plasticity, a type of developmental plasticity, is a well-documented phenomenon in larvae of the echinoid echinoderm, *Dendraster excentricus*. A recent study in our lab has shown that this morphological plasticity is associated with significant physiological plasticity for growth. The goal of the current study was to measure several aspects of protein metabolism in larvae growing at different rates to understand the mechanistic basis for this physiological growth plasticity. Larvae of *D. excentricus* were fed rations of 1,000 (low-fed) or 10,000 (high-fed) algal cells mL^−1^. Primary measurements of protein growth, algal ingestion, aerobic metabolism, alanine transport and protein synthesis were used to model growth and protein metabolism. Relative protein growth rates were 6.0 and 12.2 % day^−1^ for low- and high-fed larvae, respectively. The energetic cost of protein synthesis was similar between both treatments at 4.91 J (mg protein synthesized)^−1^. Larvae in both treatments used about 50% of their metabolic energy production to fuel protein synthesis. Mass-specific rates of protein synthesis were also similar. The most important difference between low- and high-fed larvae were mass-specific rates of protein degradation. Low-fed larvae had relatively low rates of degradation early in development that increased with larval age, surpassing high-fed degradation rates at 20 days post-fertilization. Changes in protein depositional efficiency during development were similar to those of larval growth efficiency, indicating that differences in protein metabolism are largely responsible for whole-organism growth plasticity. Mass-specific alanine transport rates were about 2-times higher in low-fed larvae, demonstrating that the longer arms of low-fed larvae may be a mechanism for acquiring more dissolved nutrients from their environment. In total, these results provide an explanation for the differences in growth efficiency between low- and high-fed larvae and demonstrate the importance of protein degradation pathways in establishing these growth differences. These observations, together with previous studies measuring morphological and physiological plastic responses, allow for a more integrated understanding of developmental plasticity in echinoid larvae.

## Introduction

The earliest stages of an organism’s life are typically characterized by both rapid development and growth. The timing, rates, and efficiencies of these processes can have important ramifications for the remainder of the organism’s life. This is especially important for marine ectotherms, whose development and growth are sensitive to many different environmental parameters. Therefore, it is critical to gain a foundational understanding of the molecular, biochemical, physiological, and morphological pathways and strategies that are utilized during these unique moments where rates of growth and development are high, relative to later stages in life. Such an understanding is critical to predicting organismal responses in the face of rapid environmental changes, many the result of anthropogenic activities.

Increase in biomass during early development, through the creation of new tissues as well as increases in biochemical reserves, occurs through accumulation of three primary biochemical substrates: proteins, lipids, and carbohydrates. While there is interspecific variability in the importance of each, it is generally observed that proteins are the most important of these biomass constituents (Fraser and Rogers, 2007). Proteins comprise the majority of an organism’s biomass and they serve as developmental and metabolic regulators because of their role as enzymes. Proteins are considered the most expensive molecules to synthesize (Hawkins, 1991; Houlihan, 1991; Berg et al., 2012), with each addition of an amino acid to a polypeptide chain costing, at a theoretical minimum, 4 ATP equivalents (Berg et al., 2012). Total costs of protein synthesis are likely much higher due to the additional costs of supporting processes such as amino acid transport, RNA synthesis, and protein trafficking. Proteins also undergo relatively high rates of turnover in which they are degraded and resynthesized, adding another substantial energetic cost due to the ATP-dependent nature of ubiquitin targeting and proteasomal degradation (reviewed in Finley, 2009). These processes are collectively classified as protein metabolism. A critical question that emerges from studies of protein metabolism and growth is whether protein growth is achieved primarily through increased synthesis rates, or by decreased degradation rates (Fraser and Rogers, 2007). Another important question is if protein growth rates can be enhanced by manipulating the efficiency of protein metabolism. For example, increased rates of protein growth could theoretically be supported by increasing protein depositional efficiency and/or decreasing the energetic cost of protein synthesis.

Much research examining growth and development during early life stages has focused on planktotrophic (feeding) marine invertebrate larvae. This is especially true for the larvae of echinoid echinoderms, where there is a relative wealth of information concerning rates of morphological growth and development and their sensitivity to critical environmental parameters (e.g., McEdward, 1984; Hart and Strathmann, 1994; Bertram and Strathmann, 1998; Reitzel et al., 2004; Byrne et al., 2008; Pan et al., 2015; Rendleman and Pace, 2018). In addition, several studies have assessed the biochemical and metabolic importance of protein metabolism during echinoid development (Marsh et al., 2001; Pace and Manahan, 2006, 2007a; Pan et al., 2015; Pan et al., 2018). These studies have clearly demonstrated that regulation of protein metabolism is a critical feature in determining organismal rates of growth and as well as an important response mechanism to environmental changes.

Echinoid larvae exhibit food-induced phenotypic plasticity (e.g., Hart and Strathmann, 1994; Miner and Vonesh, 2004; Soars et al., 2009; Adams et al., 2011). From a morphological perspective, the pluteus larva responds to different levels of unicellular algal food by changing the length of their larval arms (ciliated projections that create a feeding current that allows for the capture and ingestion of algal food) such that low-fed larvae grow longer arms to increase their feeding ability while high fed larvae grow shorter arms and allocate resources towards faster growth (Strathmann et al., 1992; Miner, 2011). This response allows high-fed larvae to attain metamorphic competency sooner, thereby reducing their time in the plankton and its associated dangers (Rumrill, 1990). It has recently been demonstrated that in conjunction with this morphological plasticity there is also significant physiological plasticity (Rendleman et al., 2018). Low-fed larvae of the echinoid *Dendraster excentricus* (Eschscholtz, 1831) have relatively (compared to high-fed larvae) high digestive efficiency that complements their increased feeding ability. This high digestive efficiency is accompanied by similarly high growth efficiencies, including protein growth efficiency (the amount of protein biomass growth relative to the amount of protein ingested). While these high efficiencies quickly diminish over development and are rapidly surpassed by high-fed larval efficiencies, they are clearly beneficial and allow the morphological and physiological plasticity responses to operate in tandem to allow the larvae to make the most of their limited food environment. While there have been no direct measurements of protein metabolism with regards to this plasticity response, some studies have indirectly provided support of differences in protein metabolism by way of thyroxine signaling (Heyland and Hodin, 2004) and target of rapamycin (TOR) signaling (Carrier et al., 2015).

The goal of the current study was to directly quantify components of protein metabolism – protein growth, synthesis, degradation, depositional efficiency, and energetic costs – in larvae of *D. excentricus* during food-induced plasticity to understand their potential roles in establishing the large differences in growth efficiency between low- and high-fed larvae. We discovered that mass-specific rates of protein synthesis, as well as costs of protein synthesis, were similar in low- and high-fed larvae. The most significant difference was in protein depositional efficiency (amount of synthesized protein retained as biomass), indicating that differential rates of protein degradation provide the mechanism by which different rates of protein growth were achieved between low- and high-fed larvae. Our results provide a physiological mechanism to explain the previously observed differences in protein growth efficiency (Rendleman et al., 2018) resulting from physiological plasticity.

## METHODS

### Sand Dollar Collection, Spawning, and Larval Culturing

*Dendraster excentricus* (Eschscholtz, 1831) adults were collected from Los Angeles Harbor in San Pedro, CA (33.7088, -118.2806). Animals were kept in large coolers during transport to the California State University, Long Beach (CSULB) Marine Laboratory. The collected sand dollars were kept in 200L tanks of flowing seawater at about 16°C for no longer than 3 days before being used for experiments. Coelomic injections of 0.5 mol L^−1^ KCl were used to induce gamete release. Sperm were diluted (1:1000) and gently mixed with eggs in sterile-filtered seawater (0.2 μm pore) until achieving a sperm to egg ratio of ~ 5:1. Fertilization envelopes were counted to confirm successful fertilization (>90%). All cultures were reared at 16 (±1)°C in the CSULB marine laboratory. For each culture, eggs and sperm from 3 females and 3 males were combined for fertilization. Embryos were reared at 5 individuals mL^−1^ in 20-L food-grade vessels (Cambro, Huntington Beach, CA). Cultures were gently mixed using a motor-driven plastic paddle (Buehler Products, NC) at 6 rpm. Experimental results (unless otherwise noted) were derived from 3 independent cultures that were initiated during July 2017 (Culture 1), February 2018 (Culture 2), and July 2018 (Culture 3).

After reaching the feeding larval stage, animals were divided equally between low- and high-fed treatments. Larvae were fed *Rhodomonas* sp., an algae commonly used to rear echinoid larvae due to its ability to support robust growth and development in a laboratory setting (e.g., Strathmann, 1971; Schiopu et al., 2006; Rendleman et al., 2018). low- and high-fed rations were fed 1,000 and 10,000 algal cells mL^−1^, respectively. Previous research (Rendleman et al., 2018) has demonstrated that these culture conditions result in the expression of both morphological and physiological plasticity and thus are appropriate for our analysis. Algae were cultured in Erlenmeyer flasks with sponge stoppers and grown using f/2 media. Algal cultures were always harvested for larval feeding at the end of their logarithmic growth phase. To minimize the potential for microbial growth and other non-specific effects related to algal nutrient media, all media was removed from algae by centrifugation (Beckman Coulter Avanti J-E : 3,000 rpm, 12 minutes, 10°C) and algae were resuspended in fresh seawater before counting and addition to larval cultures. Algal concentrations were determined using a BD Accuri C6 (BD Biosciences, San Jose, CA) flow cytometer following established methods (Cucci et al., 1989; Lizárraga et al., 2017; Rendleman et al., 2018). Algal cells were identified by chlorophyll autofluorescence stimulated by a blue laser (488 nm) and detected after passage through a 585/40 nm filter. Gating parameters and the accuracy of flow cytometer counts were established and checked using serial dilution assays of *Rhodomonas* sp. and comparison against hemocytometer counts. Algal concentrations in each larval culture were checked daily and restocked to the target feeding concentration to ensure consistent feeding conditions. High-fed larval cultures were terminated at 25 days post-fertilization (DPF) due to a large proportion of larvae metamorphosing in the culture vessels. Low-fed larval cultures were studied until 42 DPF at which point there were not enough larvae to continue the analysis. Instantaneous mortality rates (m) of all cultures were determined using the equation N_t_ = N_0_ e^−tm^ (Rumrill, 1990), where N_0_ is the number of larvae at 3 DPF, N_t_ is the number of larvae remaining at 23 DPF (a common range for both feeding treatments was used), and t is the total time, 20 days.

### Larval Morphometrics

The occurrence of morphological plasticity was confirmed by linear measurements of low- and high-fed larvae. Because it has been demonstrated that the feeding conditions used here result in morphological plasticity, the current study only examined one of the cultures (Culture 3) as a confirmation. Post oral (PO) arm length and mid-line body length (MBL) were measured as described in Rendleman et al. (2018). Measurements were made on 3, 5, 7, 10, 12, 15, and 20 DPF. Ten larvae from each treatment were removed and photographed using a QIClick digital camera mounted on an Olympus BX51 compound microscope. Lengths were then determined using Image J software calibrated using an image of a stage micrometer.

### Protein biomass growth

Throughout larval development, triplicate samples of 500 larvae were taken for measuring protein biomass. Larval concentration was estimated using a Sedgewick Rafter counting chamber. Samples of known concentration were placed in Eppendorf 1.5 mL micro centrifuge tubes. Samples were spun at 1,000g (4C) for 2 minutes and excess seawater was removed by aspiration. Samples were then stored at −80°C until protein biomass was analyzed (within 4 weeks of sample being taken). Protein biomass was determined using a BCA Protein Assay Kit (ThermoFisher, MA). Previously frozen larval samples were removed from −80°C storage and resuspended in 500 μl of NANOpure (Barnstead) water and sonicated (30% amplitutde, 3 pulses at 2 seconds each while on ice). Three aliquots of each preparation (technical replicates) were run for protein content following manufacturer’s instructions. For all determinations of protein content, known absolute amounts of Bovine Serum Albumin (BSA) were used to construct a standard curve for quantification of protein content from spectrophotometric absorption values (562nm). Protein biomass values were used to quantify biomass growth rates as well as to standardize physiological rates to biomass (e.g., respiration, amino acid transport, and protein synthesis rates). Relative growth rates (RGR, % protein mass day^−1^) were calculated using the following equation from Conceição et al. (1998):

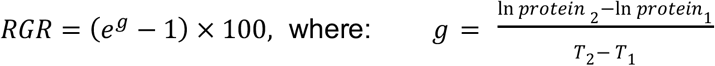

Where *e* is 2.718 (inverse of the natural log), T_1_ and T_2_ are the developmental timepoints where protein biomass samples were taken and protein_1_ and protein_2_ are the protein biomass values (in ng individual^−1^) for the respective time-points. The RGR was calculated for every interval in which protein values were taken, resulting in 6 estimates per culture for low-fed larvae and 4-5 estimates per culture for high-fed larvae.

### Ingestion Rates

Daily algal ingestion rates were measured in all 3 cultures as described in Rendleman et al. (2018). Algal concentrations were determined by measuring the decrease in algal concentration ~24 hours after restocking each larval culture vessel to its target concentration. Algal concentrations in vessels were determined using a BD Accuri^TM^ C6 flow cytometer. Three algal samples (75 μl each) were removed from each culture vessel using a 50 μm-pore Nitex mesh to ensure no larvae entered the sample. The volume of sample necessary to count 100,000 cells was recorded. Three control vessels (3 L each, being mixed with similarly proportioned paddles driven by small Beuhler motors) containing only algae were used to correct for any changes in algal concentration not attributed to larval feeding. Daily estimation also ensured that despite differences in feeding rates, low- and high-fed larvae were always restocked to their target algal concentrations. The following equation was used for ingestion rate determination:

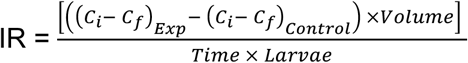

where IR = ingestion rate in the units of algal cells larva^−1^ day^−1^. Exp and Control represent concentrations for the experimental (vessels containing algae and larvae) and control (vessels only containing algae) culture vessels, respectively. C_i_ and C_f_ represent initial and final algal concentration for the experimental and control vessels in the units of algal cells mL^−1^. Volume equals the volume of the culture vessels in the units of mL. Time is the duration between the initial and final algal concentration measurements (typically between 20 to 24 hours) and larvae is the number of larvae in the experimental vessel for which ingestion rates were being estimated. The concentration of larvae in the vessels was determined 3 times per week (during water changes using a Sedgewick Rafter counting chamber) and used with culture volume to estimate larval numbers. For IR determinations on a non-water change day, larval number was estimated by interpolating from larval estimates preceding and succeeding that measurement. These IR values were then used to directly determine the amount of energy ingested through algal feeding and subsequently used to support measured physiological rates (e.g., growth, respiration, protein synthesis). Because there were instances when larvae ate most of the food provided within the 24-hour interval (especially later in development), these estimates are not reflective of their feeding capacity.

### Respiration Rates

Larval respiration rates were determined in all 3 cultures as described in Rendleman et al. (2018),using the μBOD method (Marsh and Manahan, 1999). Sterile filtered, oxygen saturated seawater was placed in 11 custom-made μBOD vials (volume ~ 800μl) and a range of larvae (50-600 individuals, depending on developmental stage) were micro-pipetted into eight μBOD vials. The additional three vials contained only filtered seawater and served as blanks to account for non-larval consumption of O_2_ (e.g., microbial respiration). The vials were then incubated at 16°C for three hours, after which time, the oxygen content was measured using a polarographic oxygen sensor (Strathkelvin Instruments, Scotland). Vials were spun down and approximately 400 μl of seawater were removed with a gas tight syringe and injected into the oxygen sensor chamber (volume ~ 70μl). After one minute, the time and oxygen concentration were recorded. The number of larvae in each vial were counted for each treatment and oxygen consumption (pmoles O_2_ hr^−1^) was linearly regressed against the number of individuals in each vial to determine pmole O^2^ hr^−1^ individual^−1^. For all respiration rates recorded, no evidence of concentration-dependent artifacts were observed (i.e., total O_2_ consumed increased linearly with increasing larvae in each μBOD).

### Efficiency Analyses

Data for daily ingestion rates, protein growth rates, and respiration rates were used to construct energetic efficiency models for assimilation efficiency (AE), gross growth efficiency (GGE), net growth efficiency (NGE), and protein growth efficiency (PGE). These values were determined following the protocols of Rendleman et al. (2018). The following equations were used for the respective efficiencies:

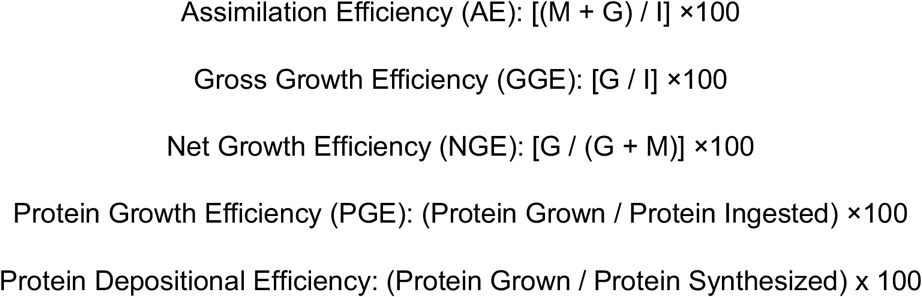

M is the energy metabolized and measured through oxygen respiration rates. Oxygen consumption values were converted to energetic units using the oxyenthalpic value of 484 kJ mol^−1^ O_2_ (Gnaiger, 1983). G is biomass growth and was determined using changes in protein biomass. Protein values were converted to energetic equivalents using a conversion of 24 kJ g^−1^ (Gnaiger, 1983; Schmidt-Nielsen, 1997). I represents energy acquired through ingestion. Ingestion rates of algal cells were converted to energetic units using the specific energetic content of *Rhodomonas* sp., 2.25 μJ (algal cell)^−1^ (Vedel and Rissgard, 1993). Protein ingested was determined using the protein content of *Rhodomonas* sp., 0.053 ng protein (algal cell)^−1^ (Rendleman et al., 2018). In order to accurately model these efficiency metrics over development, primary data (ingestion, protein growth, respiration, and protein synthesis) from all three cultures were used to create a best-fit model describing how each variable changed throughout larval development in low- and high-fed larvae. For each culture, the linear regression values (slope and y-intercept) were determined for the log_10_(x) transformed data (ingestion, respiration, or protein biomass) plotted against DPF. In order to take into account the slight variation in sample size per culture (i.e., the number of developmental time points measured), the weighted mean for each regression value was calculated (Sokal et al., 1987). These weighted average regression values were used to model changes in each respective variable throughout larval development for low- and high-fed larvae.

### Rates of protein synthesis

Rates of protein synthesis were determined as described in other studies of echinoderm larvae (Pace and Manahan, 2006; Pace et al., 2010; Pan et al., 2015). ^14^C-alanine (Perkin Elmer, MA), was used as a tracer to follow the rate of incorporation of amino acids into protein. Previous studies on echinoid larvae have shown alanine to be optimal for use as a radiolabeled tracer as it is transported rapidly from seawater into the free amino acid pool (FAA pool) and subsequently incorporated into the protein pool in sufficient amounts to be accurately measured by HPLC and liquid scintillation counting (Marsh et al., 2001; Pace and Manahan, 2006, 2007a). As part of this measurement, rates of alanine amino transport were also determined (described below). Each protein synthesis determination was a 3-part kinetic assay determining 1) rates of total alanine incorporation into larvae (i.e., alanine transport rates), 2) changes in the specific activity of alanine in the precursor free amino acid (FAA) pool, and 3) rates of alanine incorporation into larval protein. The three kinetic assays were run in parallel with each other from the same stock of larvae being exposed to the ^14^C-alanine tracer. For each experiment, a known number of larvae were exposed to 10-20 μmol L^−1^ ^14^C-alanine in 10ml of seawater (total radioactivity = 5μCi). Each assay was composed of 6 individual timepoints, taken over the course of 20-25 minutes, used for calculating linear rates of change for each respective variable. The short time frame minimizes the effect of protein degradation, where ^14^C-alanine reenters the free amino acid pool due to protein degradation. The samples of larvae were placed onto an 8 μm filter (Nucleopore) and washed with 3-times with FSW to remove unincorporated radioactivity. Using these data, whole animal rates of protein synthesis were determined using the following equation:

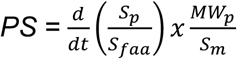

Where PS = protein synthesis rate in ng larva^−1^ hr^−1^, S_p_ = radioactivity in the total protein pool (procedure described below), S_faa_ = specific activity of the free amino acid pool (procedure described below), MW_p_ = average molecular weight of an amino acid in *D. excentricus,* and S_m_ = the mole percent of the amino acid used in protein. The following descriptions detail the methods for acquiring the different data sets represented in the above protein synthesis equation.

#### Specific activity of the free amino acid pool (S_faa_)

The S_faa_ was determined through high performance liquid chromatography (HPLC) separation of the free amino acid pool (FAA) and liquid scintillation counting (Beckman Coulter LS 6500, Brea CA) of the alanine elution peak. A known number of larvae were exposed to 10-20 μmol L^−1^ ^14^C-alanine in 10ml of seawater (total radioactivity = 5μCi). Six 500 μl samples were taken regularly over a 25-minute interval to quantify the temporal increase in the S_faa_. The FAA for each sample was extracted in 70% ethanol and alanine was separated from other amino acids using reverse phase HPLC (Lindroth and Mopper, 1979). Amino acids were detected using precolumn derivatization with ortho-phthalaldehyde (OPA) (Sigma Aldrich P0657). A dual gradient solvent system in conjunction with a C-18 reverse phase column separated amino acids based on hydrophobicity. The gradient system progressed from hydrophilic (Solvent A: 80% 50mM sodium acetate, 20% methanol) to hydrophobic (Solvent B: 20% 50mM sodium acetate, 80% methanol). Alanine was quantified through peak area comparison with standards of known concentrations. The alanine elution peak was collected in a fraction collector and its radioactivity was determined by liquid scintillation counting. This method allowed for accurate determination of changes in the precursor specific activity in the units of dpm pmol^−1^ alanine. This value was then used to correct for the radioactive incorporation rate of ^14^C-alanine into the protein pool.

#### Radioactivity in the total protein pool (S_p_)

Larvae sampled for determining the rates of ^14^C-alanine incorporation into protein were immediately frozen in liquid nitrogen. For each sample 500 larvae were collected. These samples were subsequently sonicated and total protein was precipitated using cold 5% trichloroacetic acid (TCA). The protein precipitate was collected on a GF/C glass microfiber filter (Whatman, Sigma Aldrich) and then mixed with EcoScint (Fisher Scientific) scintillation cocktail. Radioactivity of the protein was determined using a Beckman (LS 6500) liquid scintillation counter. The amount of ^14^C-alanine incorporated into total protein was converted to total alanine incorporation using the measured intracellular specific activity (^14^C per mol total alanine in the free amino acid pool, aka S_faa_). By standardizing the S_p_ by the S_faa_ over the duration of the experiment, the rate of total alanine incorporation was determined (pmol alanine larva^−1^ hr^−1^).

#### Average molecular weight of an amino acid in *D. excentricus* and mole-percent alanine in total protein (MW_p_ and S_m_)

In order to convert alanine incorporation rates into total amino acid incorporation rates (i.e., rate of protein synthesis), the amino acid composition of larval protein (i.e., mole percent composition of protein; S_m_) and the average molecular weight of an amino acid in the protein pool (MW_p_) were determined. Amino acid composition was determined by UC Davis Core Facility using an amino acid analyzer. Protein was extracted in 5% TCA and then acid-hydrolyzed in 6 mol l^−1^ HCl with 0.5% phenol at 110°C for 24 hours under vacuum. The acid-hydrolyzed protein was then run on a cation exchange chromatographer. Post-column ninhydrin derivitization was used to find amino acid composition.

#### Total alanine larval transport

Alanine transport rates were determined by measuring the increase in total larval radioactivity. While transport rates are not part of the protein synthesis calculation, they were conducted in conjunction with protein synthesis experiments. This data was also us to construct radioactivity budgets to ensure the sum of ^14^C alanine incorporation into the FAA and the total protein pool were equal to the total radioactive incorporation measured as alanine transport. Larvae were sampled and placed in a liquid scintillation vial with 5 ml of EcoScint cocktail (Fisher Scientific). Samples were counted in a Beckman (Brea, CA) LS 6500 scintillation counter. To ensure maximal rates of transport, ^12^C-alanine (i.e., “cold” alanine) was added to the experimental vial to bring the final concentration to 20μM. Alanine transport rates were then corrected using the necessary ^14^C:^12^C-alanine ratio (hot:cold ratio). Alanine transport rates were also determined in experiments using the protein synthesis inhibitor, anisomycin, in order to assess if the inhibitor had any non-specific effects.

### Cost of Protein Synthesis

The energetic cost of protein synthesis [J (mg protein synthesized)^−1^] was calculated using two different approaches. Both approaches utilized respiration rate and protein synthesis rate data, acquired as described above. A direct approach used inhibition of protein synthesis to calculate its cost at specific developmental time-points (i.e., prefeeding larvae as well as in low- and high-fed larvae at 11 and 19 DPF). This approach measured the change in respiration and protein synthesis when in the presence or absence of the eukaryotic protein synthesis inhibitor, anisomycin (Sigma-Aldrich, St. Louis, MO), which has been shown to be highly effective and specific (Fenteany et al., 1995; Pace and Manahan, 2007a). Anisomycin was used at a concentration of 10μM. For calculating the cost of protein synthesis, rates of oxygen consumption were converted to energetic units using an oxyenthalpic conversion of 484 kJ mol^−1^ O_2_ (Gnaiger, 1983). Costs were determined by dividing the difference in metabolic rate by the difference in protein synthesis rates when in presence of the inhibitor. Therefore, the efficacy of the inhibitor was never assumed but empirically determined. Additionally, rates of alanine transport were measured during inhibitor exposure as a check for potential non-specific effects of the inhibitor.

We also used an indirect, correlative approach to determine the cost of protein synthesis throughout larval development. Here the relationship between changes in respiration rate and protein synthesis rate over development was assessed. These values represent normal physiological rates (i.e., no inhibitors were employed). Respiration rates were plotted (ordinate) against protein synthesis rates (abscissa) and tested for a significant relationship. The slope of this relationship therefore represents the cost of protein synthesis (the change in energy consumption rate per unit increase in protein synthesis rate).

### Rates and costs of protein degradation

Rates of protein degradation were calculated as the difference between the modeled rates of protein synthesis and protein growth. The cost of protein degradation was taken from Pan et al. (2018) as 0.14 kJ gram^−1^ protein degraded. As specified in Pan et al. (2018), this cost is based on the primary analysis by Peth et al. (2013) where degradation is by the 26S proteasomal pathway. This estimate is based on the assumptions that degradation occurs via the 26S pathway and requires 0.003 mol ATP gram^−1^ protein and that 45 kJ of energy are liberated with the hydrolysis of a mole of ATP [details in Pan et al. (2018)].

### Statistical Analysis

Variation in morphological and physiological variables as a function of developmental time (DPF) and feeding treatment (low vs. high) were evaluated with general linear models (GLMs). Replicate cultures were treated as a random factor so that within treatment variation among cultures could be assessed. In the absence of a statistically significant interaction between main effects, analysis of covariance (ANCOVA) was used to compare adjusted mean values of the different feeding treatments; when there was a significant interaction, no further analysis was done. Prior to analyses, physiological variables were transformed using a log_10_(x) function to correct for non-linearity, non-normality, and unequal variances as necessary. Visual inspection of model residuals was done for every analysis.

Regression slopes and intercepts for the three independent cultures were pooled within feeding treatments as weighted mean averages and weighted mean standard deviations to parameterize models of rate changes during development (after Sokal et al., 1987). In cases where there was a significant block effect (indicative of culture to culture variation), data were still averaged because the main effects were always much larger than the block effect (as determined by Eta squared, the ratio of effect sum of squares to total sum of squares (Thompson, 2006)) and the general outcome remained the same. All analyses were performed using Minitab 18.

## Results

### Development and Induction of Morphological Plasticity

Both feeding treatments were effective in supporting larval growth and development (Fig. 1A). Average rates of mortality for the three cultures were similar between feeding treatments (ANOVA, *F*_*1,5*_ = 7.71; *P* = 0.57) at 0.067 ± 0.023 day^−1^ and 0.056 ± 0.023 day^−1^ (mean ± SD, N=3) for low- and high-fed larvae, respectively. High-fed larvae reached metamorphic competency by at least 24 DPF, where it was observed that they were metamorphosizing in the culture vessels. Low-fed larvae were not observed to reach metamorphosis in any of the culture vessels.

**Figure 1.**
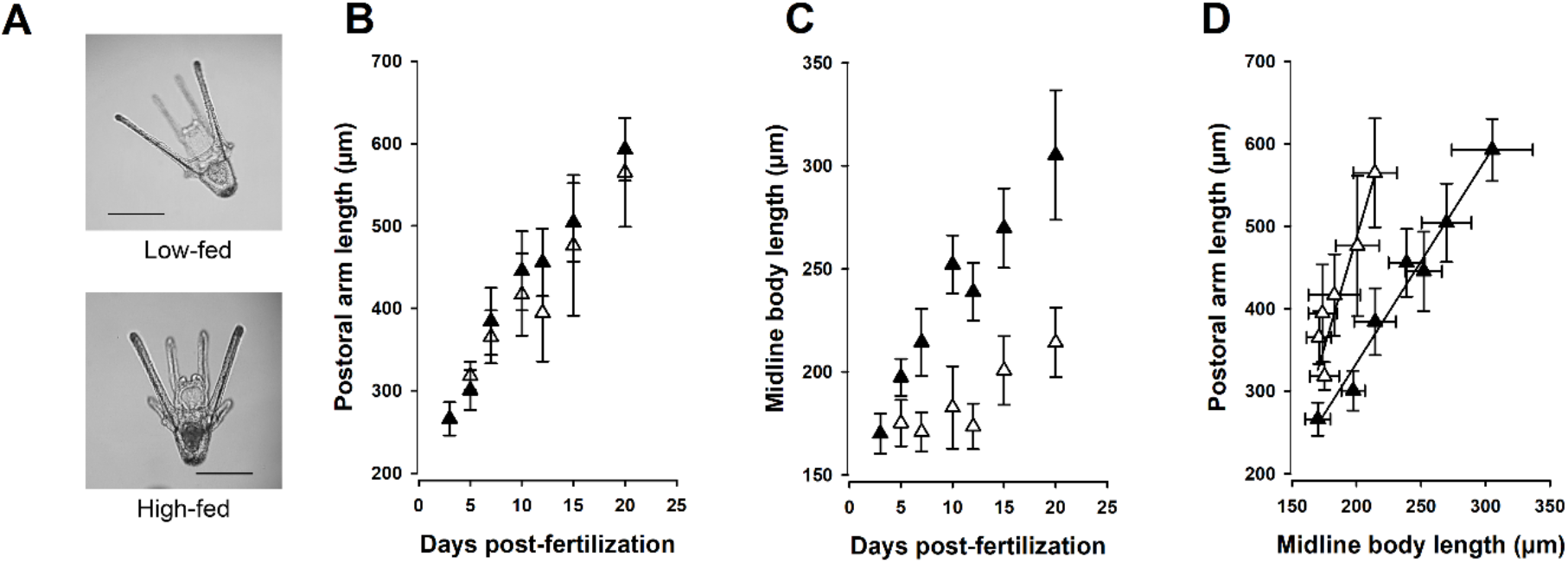
Changes in larval morphology and expression of morphological plasticity in low- and high-fed larvae of *D. excentricus*. **(A)** Larvae at 7 DPF. Scale bar = 200μm. **(B)** Postoral arm growth in low- and high-fed larvae. There was no significant treatment effect (ANOVA, *F*_*1,13*_ = 0.04, *P* = 0.85) or interaction with larval age (ANOVA, *F*_*1,13*_ = 1.44, *P* = 0.26), indicative that PO arm length growth was similar in low- and high-fed larvae. **(C)** Midline body length growth in low- and high-fed larvae. **(D)** Postoral arm growth as a function of midline body length. Solid line represents linear regression model for each feeding treatment. Low-fed: y = 5.34x – 581.0, r^2^ = 0.84; High-fed: y = 2.47x – 159.3, r^2^ = 0.98. There was a significant feeding treatment effect: ANOVA, *F*_1, 13_ = 11.36, *P* = 0.007, indicative of low-fed larvae having a longer postoral arm length for any given midline body length. For all graphs: low-fed = white symbols; high-fed = black symbols. Data for morphology was from Culture 3.

Despite the order of magnitude difference in available food, postoral (PO) arm growth was similar between low- and high-fed larvae (ANOVA, *F*_*1,13*_ = 0.04, *P* = 0.85) (Fig. 1A and Fig. 1B) and there was no interaction between feeding treatment and larval age (ANOVA, *F*_*1,13*_ = 1.44, *P* = 0.26). In contrast, for midline body length (MBL) a significant interaction between feeding treatment and age was observed (ANOVA, *F*_*1,13*_ = 27.76, *P* < 0.001), indicative of faster rates of MBL growth in high-fed larvae (Fig. 1C). The relationship between PO length and MBL exhibited a significant feeding treatment effect (ANOVA, *F*_*1,13*_ = 11.36, *P* = 0.007), indicating that at any given MBL, low-fed larvae had a longer PO length (Fig. 1D). These results are similar to those in our previous study (Rendleman et al., 2018).

### Physiological Plasticity

Rates of protein growth (Fig. 2A), algal ingestion (Fig. 2B), and respiration (Fig. 2C) were measured so as to ensure the patterns of physiological plasticity that were documented in Rendleman et al. (2018) were observed in this study as well, which used identical culturing and treatment conditions. Larval protein biomass increased throughout development for both feeding treatments (Fig. 2A). There was a strong interaction between feeding treatment and larval age (DPF) (ANOVA, *F*_*1, 38*_ = 20.35; *P* < 0.001; Table 1), indicative of protein biomass starting from a common value and high-fed larvae growing at a faster rate than low-fed larvae. By 23 DPF, low-fed larvae had 309 ng larva^−1^, while high-fed larvae had 1420 ng larva^−1^ (using the common regression lines illustrated in Fig. 2A). At 42 DPF, low-fed larvae had 763 ng protein larva^−1^. Relative growth rates (RGR) were significantly different in low- and high-fed larvae (ANOVA, *F*_*1, 5*_ = 13.8; *P* < 0.05). High-fed rates were 2-times faster (Fig. 2A, inset), with average values of 6.0 (± 0.59)% day^−1^ and 12.2 (± 1.6)% day^−1^ (N=3, ± SE) for low- and high-fed larvae, respectively.

**Table 1.** Statistical analyses for physiological variables in *D. excentricus* larvae reared at d high food concentrations. Cultures were treated as a random blocking factor.

| **Trait** | **Source** | **df** | **MS** | **F** | **P** |
| --- | --- | --- | --- | --- | --- |
| **Protein biomass**<br>**(ng protein ind<sup>-1</sup>)**<br>**Fig. 2A** | Culture | 2 | 0.021 | 0.93 | 0.40 |
|  | Treatment (Low- and High-fed) | 1 | 0.093 | 4.04 | 0.052 |
|  | DPF | 1 | 5.38 | 238.87 | < 0.001 |
|  | Interaction (Treatment x DPF) | 1 | 0.11 | 20.35 | < 0.001 |
|  | Error | 38 | 0.023 |  |  |
| **Ingestion rate**<br>**(algal cells**<br>**day<sup>-1</sup> ind<sup>-1</sup>)**<br>**Fig. 2B** | Culture | 2 | 2.35 | 52.86 | < 0.001 |
|  | Treatment (Low- and High-fed) | 1 | 6.14 | 138.19 | < 0.001 |
|  | DPF | 1 | 8.23 | 185.31 | < 0.001 |
|  | Interaction (Treatment x DPF) | 1 | 0.095 | 2.14 | 0.146 |
|  | Error | 151 | 0.044 |  |  |
| **Respiration rate**<br>**(pmol O<sub>2</sub> hr<sup>-1</sup> ind<sup>-1</sup>)**<br>**Fig. 2C** | Culture | 2 | 0.34 | 23.41 | < 0.001 |
|  | Treatment (Low- and High-fed) | 1 | 0.42 | 28.59 | < 0.001 |
|  | DPF | 1 | 2.99 | 203.55 | < 0.001 |
|  | Interaction (Treatment x DPF) | 1 | 0.14 | 9.51 | 0.004 |
|  | Error | 32 | 0.015 |  |  |
| **Transport rate**<br>**(pmol hr<sup>-1</sup> ind<sup>-1</sup>)**<br>**Fig. 3A** | Culture | 2 | 381.23 | 14.75 | < 0.001 |
|  | Treatment (Low- and High-fed) | 1 | 1.76 | 0.07 | 0.796 |
|  | DPF | 1 | 3.37x10<sup>3</sup> | 130.39 | < 0.001 |
|  | Interaction (Treatment x DPF) | 1 | 557 | 21.55 | < 0.001 |
|  | Error | 32 | 25.84 |  |  |
| **Transport rate**<br>**(pmol hr<sup>-1</sup>**<br>**µg<sup>-1</sup> protein)**<br>**Fig. 3B** | Culture | 2 | 1.7x10<sup>-3</sup> | 2.83 | 0.074 |
|  | Treatment (Low- and High-fed) | 1 | 2.7x10<sup>-3</sup> | 4.49 | 0.042 |
|  | DPF | 1 | 2.3x10<sup>-3</sup> | 3.92 | 0.056 |
|  | Interaction (Treatment x DPF) | 1 | 5.0x10<sup>-6</sup> | 0.01 | 0.931 |
|  | Error | 32 | 6.0x10<sup>-4</sup> |  |  |
| **Protein synthesis**<br>**rate**<br>**(ng protein hr<sup>-1</sup>**<br>**ind<sup>-1</sup>)**<br>**Fig. 4A** | Culture | 2 | 0.047 | 0.94 | 0.401 |
|  | Treatment (Low- and High-fed) | 1 | 0.63 | 12.61 | 0.001 |
|  | DPF | 1 | 3.83 | 76.87 | < 0.001 |
|  | Interaction (Treatment x DPF) | 1 | 0.055 | 1.10 | 0.302 |
|  | Error | 32 | 0.050 |  |  |
| **Protein synthesis**<br>**rate**<br>**(ng protein**<br>**synthesized ng<sup>-1</sup>**<br>**protein biomass)**<br>**Fig. 4B** | Culture | 2 | 0.068 | 0.90 | 0.417 |
|  | Treatment (Low- and High-fed) | 1 | 0.12 | 1.59 | 0.218 |
|  | DPF | 1 | 0.44 | 5.79 | 0.022 |
|  | Interaction (Treatment x DPF) | 1 | 8.0x10<sup>-3</sup> | 0.11 | 0.747 |
|  | Error | 32 | 0.075 |  |  |
| **Correlative cost of**<br>**protein synthesis**<br>**[J (mg protein**<br>**synthesized)<sup>-1</sup> ng**<br>**Fig. 5B** | Culture | 2 | 266 | 1.29 | 0.289 |
|  | Treatment (Low- and High-fed) | 1 | 1.09x10<sup>3</sup> | 5.29 | 0.028 |
|  | Protein synthesized | 1 | 2.48x10<sup>4</sup> | 120 | < 0.001 |
|  | Interaction (Treatment x DPF) | 1 | 445 | 2.16 | 0.152 |
|  | Error | 32 | 206 |  |  |

**Figure 2.**
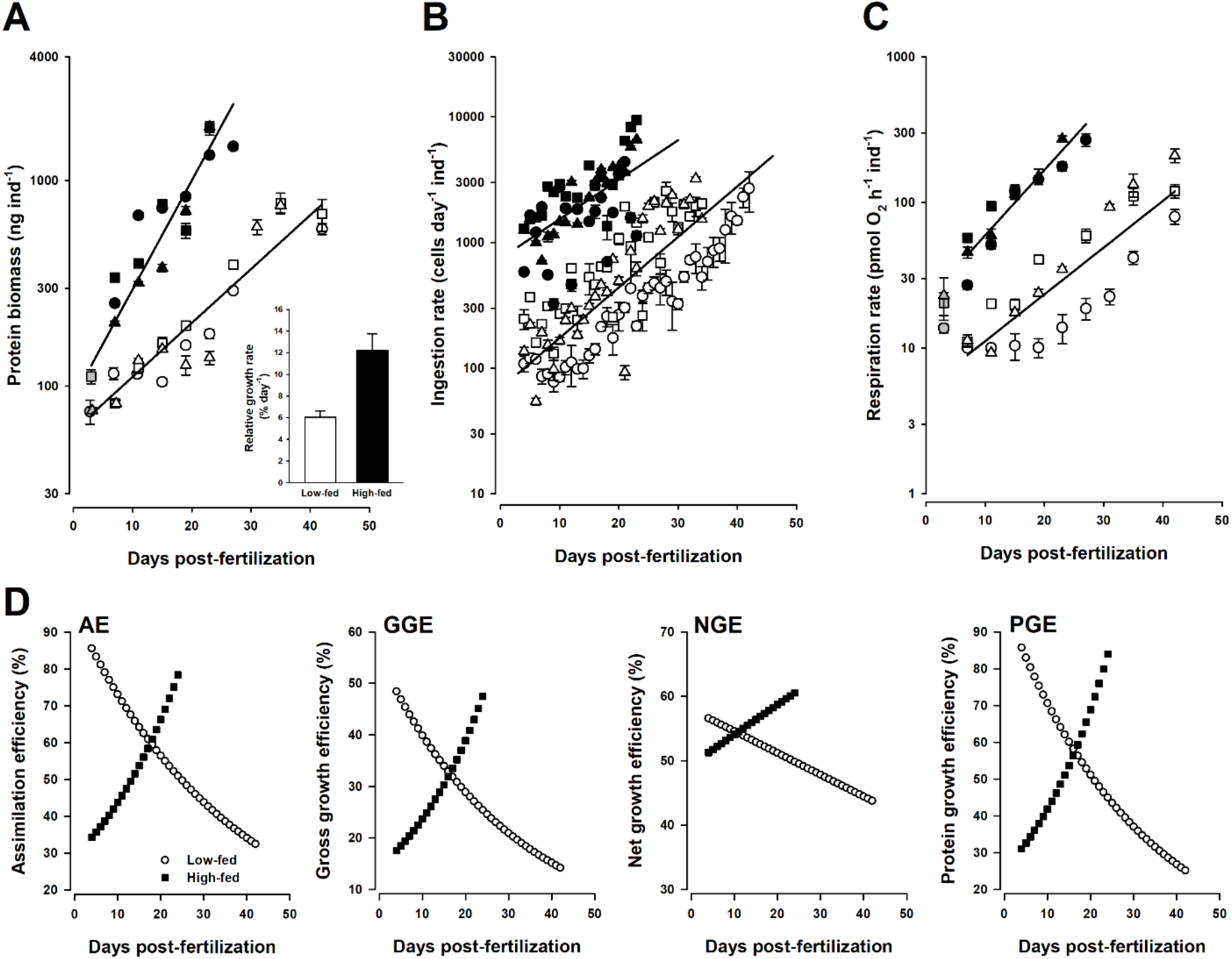
Physiological parameters and energetic efficiencies of digestion and growth in low- and high-fed larvae of *D. excentricus*. **(A)** Changes in protein biomass (plotted on log_10_ scale) in prefeeding (gray symbols), low-fed (white symbols), and high-fed (black symbols) larvae as a function of age (DFP). Symbols are means ± SEM (n = 3). Regression lines show the pooled response for low- and high-fed larvae for all three cultures analyzed: Culture 1 = circles, Culture 2 = squares, Culture 3 = triangles. Inset: relative growth rates for each treatment. Mean RGR ± SEM (n = 3). **(B)** Ingestion rates (plotted on a log_10_ scale) in low- and high fed larvae as a function of age. Symbols are means (n = 3) and are the same as in panel A. Regression lines are for all 3 cultures pooled for each feeding treatment. **(C)** Respiration rates (plotted on a log_10_ scale) in prefeeding (gray symbols), low-fed (white symbols), and high-fed (black symbols) larvae. Symbols represent respiration rate ± SEM (n = 7). Regression lines are for all 3 cultures pooled for each feeding treatment (low- and high-fed). Symbols are the same as in panel A. **(D)** Modeled daily efficiency measurements relating to digestion and growth in low- and high-fed larvae. AE = assimilation efficiency, GGE = gross growth efficiency, NGE = net growth efficiency, PGE = protein growth efficiency. All values were determined using parameters from regression lines in panels A-C. See supplementary information relating to regression values used for energetic modeling.

Rates of algal ingestion (Fig. 2B) increased for low- and high-fed larvae with development. Rates were always higher in high-fed larvae and no statistical interaction was observed between feeding treatment and larval age (Table 1). Differences in feeding rates were reflective of the 10-fold difference in available algae. For example, at 25 DPF rates of ingestion were 691 and 5,230 algal cells day^−1^ ind^−1^ for low- and high-fed larvae, respectively. Larval rates of respiration (Fig. 2C) increased significantly with larval age (*F*_*1, 32*_ = 204; *P* < 0.001; Table 1). There was a significant interaction between feeding treatment and larval age (*F*_*1, 32*_ = 9.51; *P* = 0.004; Table 1), indicating that rates of respiration were increasing faster in high-fed relative to low-fed larvae during development.

Rates of protein growth, algal ingestion, and respiration were used to model physiological efficiencies relating to digestion and growth. The regression variables to model these changes in energy acquisition were determined by taking the weighted average regression variables for each metric (taking into consideration unequal samples sizes for each independent culture). These values and their standard deviations are given in Fig. S1. Using these values, the daily changes in energy acquired, respired, and invested into growth were determined for low- and high-fed larvae (Fig. S2). These values were then used to construct the models for daily changes in assimilation efficiency (AE), gross growth efficiency (GGE), net growth efficiency (NGE), and protein growth efficiency (PGE) (see methods for details of calculations) and are displayed in Fig. 2D. For all efficiencies that were standardized to the amount of energy acquired (i.e., AE, GGE, and PGE), a common pattern was observed. Low-fed larvae initially had high efficiencies that decreased rapidly as development proceeded. High-fed larvae had low efficiencies that increased rapidly during development. This resulted in high-fed larvae efficiencies surpassing low-fed efficiencies by 18 DPF. A comparison of these efficiencies at 24 DPF (the end of the high-fed culture) shows that high-fed larvae had a 1.6-times higher AE, and 2.0 times higher GGE and PGE than low-fed larvae. Net growth efficiencies also exhibited the same diverging pattern however both low- and high-fed started at a similar value of ~ 53%, which increased to 61% in high-fed and decreased to 49% in low-fed by 24 DPF. After 24 DPF, low-fed values continued to decrease for all efficiencies metrics with the following values at 42 DPF (the last day of monitoring for low-fed larvae): AE = 32%, GGE = 14%, NGE = 44%, PGE = 25%.

### Protein Metabolism

#### Alanine Transport Rates

Rates of alanine transport increased with development for both low- and high-fed larvae (Fig. 3A). While rates were similar in early development, the increase in transport rates was faster in high-fed larvae as evidenced by a significant interaction between feeding treatment and larval age (ANOVA, *F*_*1,32*_ = 21.55, *P* < 0.001, Table 1). When transport rates were standardized to protein biomass (Fig. 3B), the developmental increase in alanine transport rates was only marginally significant (ANOVA, *F*_*1,32*_ = 3.92, *P* = 0.06, Table 1). In contrast to whole larval rates, mass-specific transport rates in low-fed were now significantly higher than rates in high-fed larvae (ANOVA, *F*_*1,32*_ = 4.49, *P* = 0.042, Table 1). Adjusted mean mass-specific transport rates (inset, Fig. 3B) were ~ 2-times higher for low-fed larvae compared to high-fed larvae (i.e., 0.088 and 0.043 pmol hr^−1^ (μg protein)^−1^, respectively).

**Figure 3.**
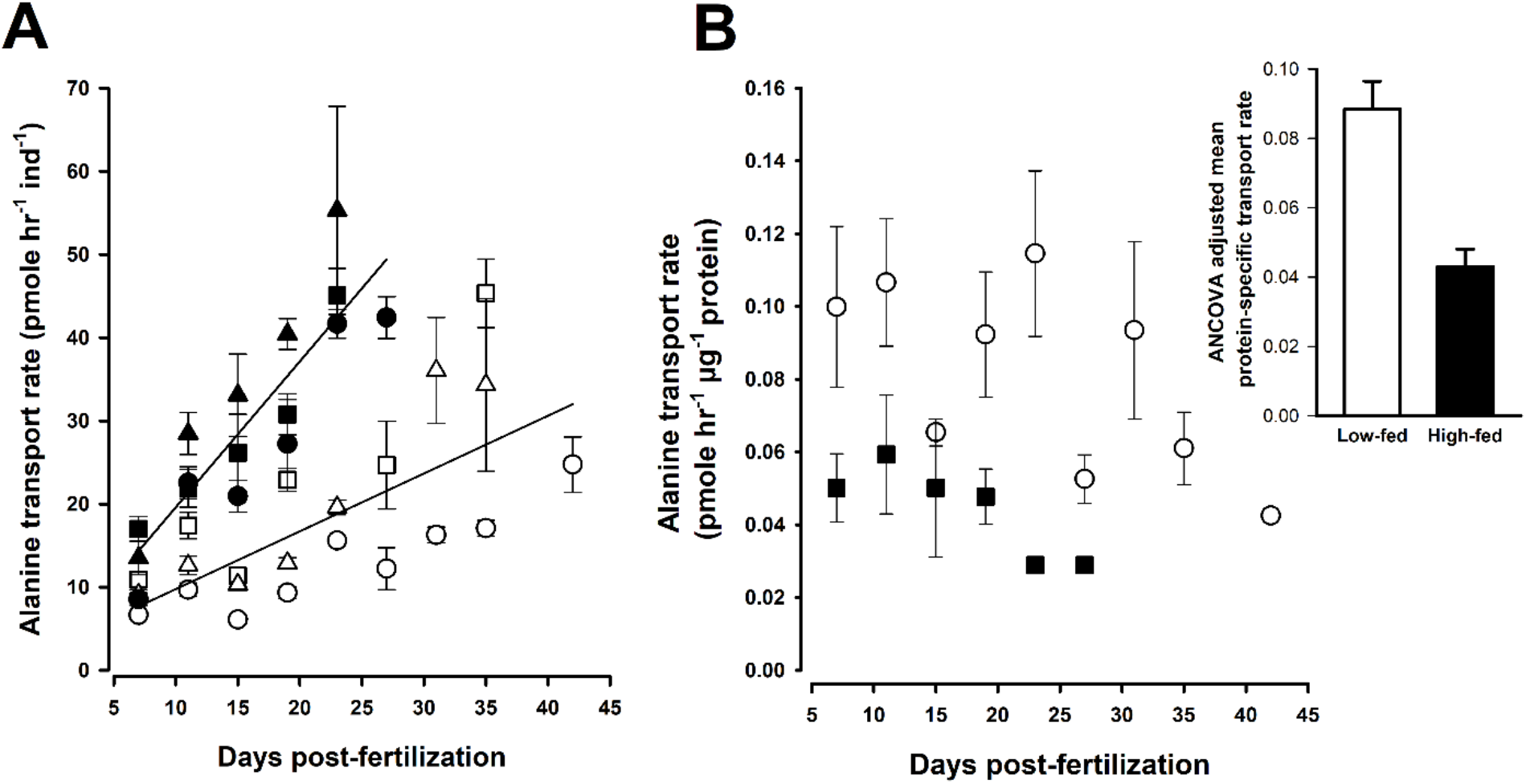
Alanine transport rates in low- and high-fed larvae of *D. excentricus.* **(A)** Whole-animal rates of alanine transport as a function of larval age for low-fed (white symbols) and high-fed (black symbols) larvae for all three cultures analyzed: Culture 1 = circles, Culture 2 = squares, Culture 3 = triangles. Regression line is for all three cultures for each treatment. **(B)** Protein-specific alanine transport rates for low-fed (white circles) and high-fed (black squares) larvae. Each value is the average transport rate (± SEM) at each larval age for the 3 replicate cultures shown in panel (A). Protein values taken from data in Fig. 2A. Inset shows the least squares means (ANCOVA) of protein-specific transport rate (from Fig. 3B), error bars are SEM. Statistical analyses for larval and mass-specific transport rates are given in Table 1.

#### Protein Synthesis Rates

Amino acid composition (both S_m_ and MW_p_) was similar for prefeeding (3 DPF) and low- and high-fed larvae (19 DPF) (Supplemental Table S1). The average percent of alanine in total protein was 9.5% ± 0.61 (mean ± SD; n = 3). The average molecular weight of an amino acid in the total protein pool (MW_p_) was 126.0 g mol^−1^ ± 0.25 (mean ± SD; n = 3).

Prefeeding rates of protein synthesis (3 DPF) were ~ 1 ng hr^−1^ ind^−1^ (Fig. 4A). Upon initiation of feeding, low-fed rates initially decreased while high-fed rates increased. As development progressed, both low- and high-fed larvae exhibited significant increases in rates of protein synthesis (ANOVA, *F*_*1,32*_ = 76.87, *P* < 0.001, Table 1). As no significant interaction was observed between feeding treatment and larval age, rates increased similarly (Table 1). However, for any given age high-fed larvae had a significantly higher protein synthesis rate than low-fed larvae (ANOVA, *F*_*1,32*_ = 12.61, *P* = 0.001, Table 1). When rates of synthesis were standardized to larval biomass (i.e., fractional rate of protein synthesis), representing the percent of the total protein pool that was being synthesized, values ranged from 0.5 to 1.5% hr^−1^ (Fig. 4B). Unlike whole organismal rates of protein synthesis, fractional rates of protein synthesis were similar between low- and high-fed larvae (ANOVA, *F*_*1,32*_ = 1.59, *P* = 0.218, Table 1), indicating that increases in whole-larval protein synthesis rates (Fig. 4A) were driven by increases in protein biomass (Fig. 2A).

**Figure 4.**
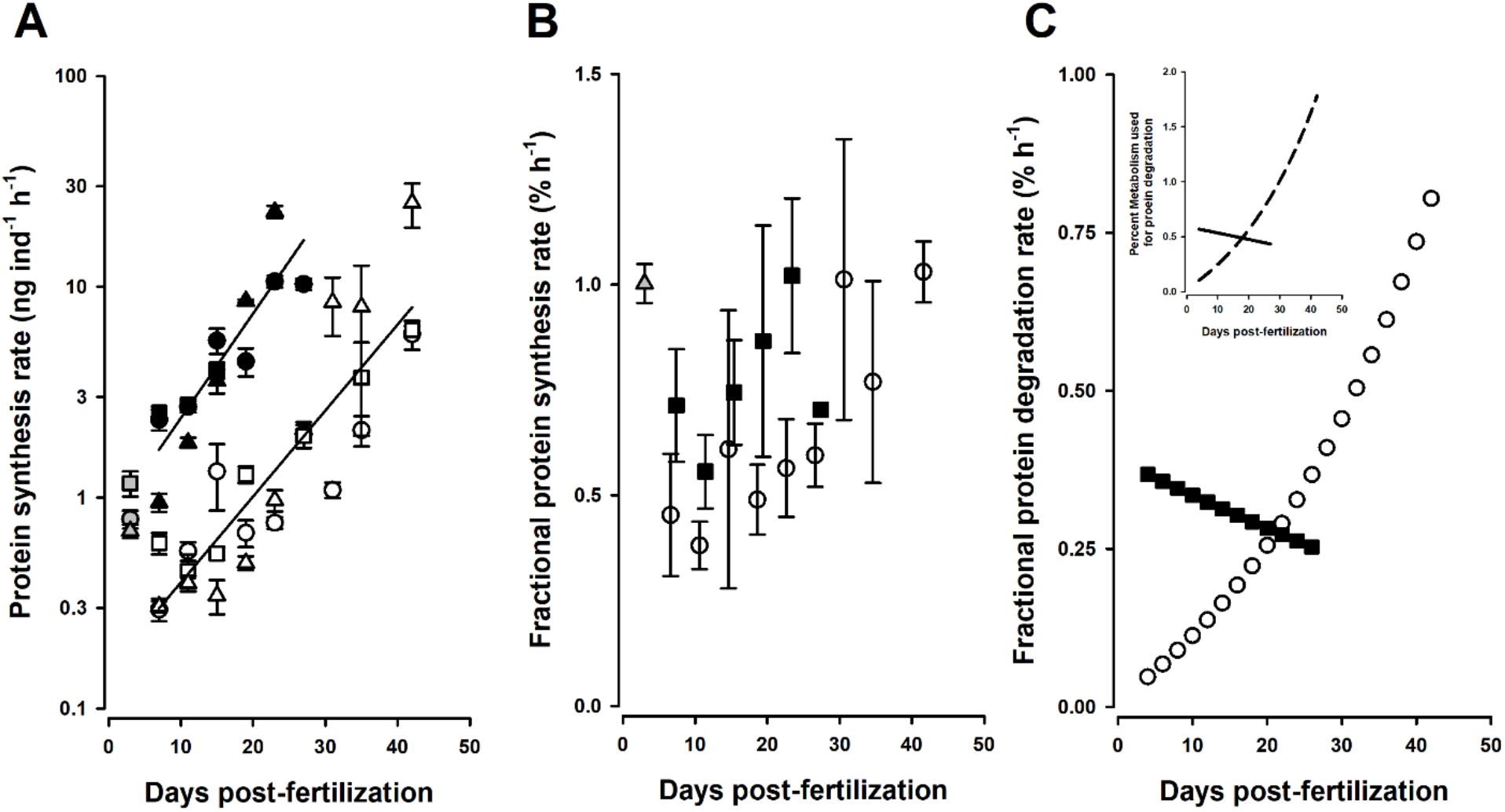
Protein metabolism in low- and high-fed larvae of *D. excentricus.* **(A)** Whole-animal rates of protein synthesis (plotted on a log_10_ scale) in prefeeding (gray symbols), low-fed (white symbols), and high-fed (black symbols) larvae as a function of age (DFP). Regression lines show the pooled response for low- and high-fed larvae for all three cultures analyzed: Culture 1 = circles, Culture 2 = squares, Culture 3 = triangles. Each data point is the result of a 6-point kinetic assay, error bar = SE of slope (see Methods for details). **(B)** Fractional rates of protein synthesis. Values represent the average for the 3 independent cultures at each larval age (+/− SEM). Values slightly offset for visual clarity. Gray triangle = prefeeding, white circles = low-fed, black squares = high-fed. **(C)** Modeled rates of protein degradation in low-fed (white circles) and high-fed (black squares) larvae. Degradation rates were estimated using the difference between protein synthesized per day (from Fig. 4A) and protein grown per day (from Fig. 2A). Inset shows the estimated percent of metabolism used for protein degradation during development. The cost of protein degradation of 0.14 kJ gram^−1^ protein was taken Pan et al., 2018.

#### Rates and metabolic importance of protein degradation

Fractional rates of protein degradation (Fig. 4C) were modeled using the data for protein growth (Fig. 2A) and protein synthesis (Fig. 4A). This assumes that for growing organisms (experiencing an increase in protein biomass) the amount of protein degradation is equal to the difference between protein synthesized and protein grown (Hawkins, 1991; Houlihan, 1991). Unlike fractional rates of protein synthesis, there was a noticeable influence of feeding treatment on rates of protein degradation. Fractional rates of protein degradation for low-fed larvae increased 17-fold (0.048 to 0.80 ng degraded ng^−1^ protein biomass) from 4-42 DPF. For high-fed, rates decreased moderately from 0.37 to 0.25 ng degraded ng^−1^ protein biomass (from 4-24 DPF). The percent of metabolism used to fuel these rates of degradation (Fig. 4C, inset) was low for both treatments, but increased for low-fed from 0.1 to 1.8% of metabolism. High-fed larvae used about 0.5% of metabolism to fuel protein degradation.

#### Energetic Cost of Protein Synthesis

The energetic cost of protein synthesis was measured using two different techniques. The first approach (Fig. 5A) used parallel measurements of protein synthesis and respiration rates with and without the protein synthetic inhibitor, anisomycin. This measurement allowed for determinations of cost at specific developmental time points. The rates used to calculate these costs are shown in Fig. 5A, with the total height of bars representing non-inhibited rates and black bars showing rates during anisomycin inhibition. The inset shows the average costs for each treatment and for prefeeding stages (3 DPF). No significant differences were observed for this direct analysis of protein synthesis cost (ANOVA, *F*_*1,5*_ = 0.82, *P* = 0.52) between prefeeding, low-fed, and high-fed larvae. The average cost for all treatments (Fig. 5A, inset) was 4.47 ± 0.63 J (mg protein synthesized)^−1^ (± SE, N = 6). To further assess the specificity of anisomycin inhibition, rates of alanine transport both with and without inhibitor were measured at 11, 19, and 23 DPF in low- and high-fed larvae (Fig. S3). There were no significant differences in transport rates between anisomycin-exposed and non-exposed larvae (paired t-test: t(5) = 0.96, *P* = 0.38).

**Figure 5.**
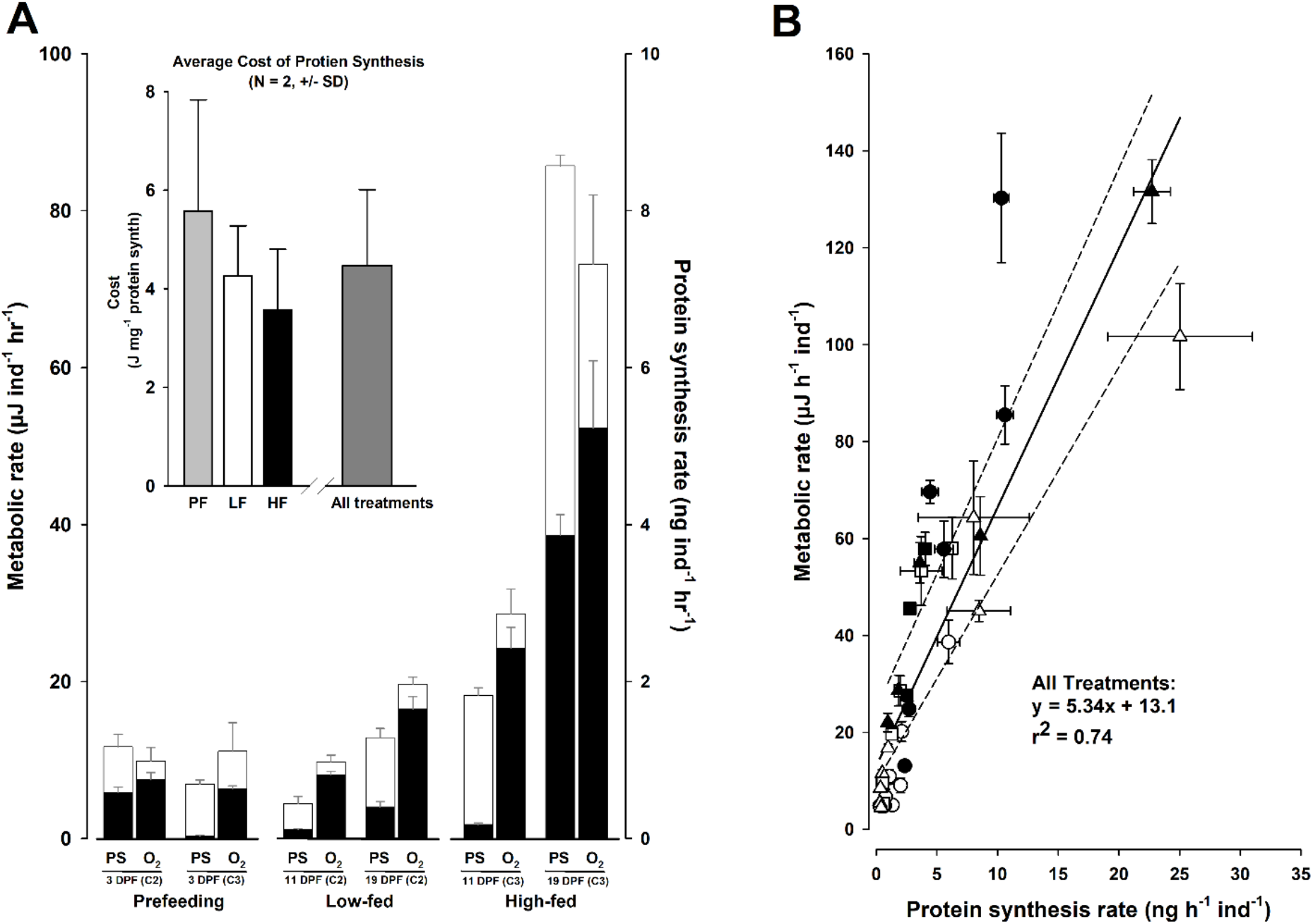
Energetic cost of protein synthesis and percent metabolism in low- and high-fed larvae of *D. excentricus*. **(A)** Primary data for inhibitor analysis calculating the energetic cost of protein synthesis. Rates of protein synthesis (PS, right y axis) and metabolism (O_2_, left y axis) for the developmental stages and feeding treatments used to calculate cost. Metabolic rates were converted to energetic units using an oxyenthalpic value of 484 Joules per mole O_2_ consumed. Total height of white bars are rates of protein synthesis and metabolism without protein synthesis inhibitor, anisomycin. Black bars are respective rates when in the presence of anisomycin. Inset shows the average cost for prefeeding stages, low-fed, and high-fed larvae derived from primary data. Differences between stage/treatments were not significant (ANOVA: df = 1, 5, P = 0.62) and the average cost was 4.47 J (mg protein)^−1^ +/− 0.63 (N=6, +/− SE). **(B)** Data for calculating the correlative cost of protein synthesis. Metabolic rates were plotted as a function of protein synthesis rates throughout development (no inhibitor was used for this approach). The solid regression line represents pooled relationship for low- and high-fed treatments: low-fed (white) and high-fed (black). Symbols represent the 3 different cultures followed in this study: Culture 1 = circles, Culture 2 = squares, Culture 3 = triangles. The slope of the relationship represents the cost of protein synthesis of 5.34 J (mg protein)^−1^ +/− 0.51 (N = 33, +/− SE). The dashed lines above and below the solid regression line represent the high-fed (above) and low-fed (below) correlative costs. As described in text, while the elevation of these correlations were different, the slope values were similar to each other.

The second approach for determining cost of protein synthesis was to investigate the average cost throughout low- and high-fed larval development by quantifying the change in metabolic rates with protein synthesis rates throughout development (Fig. 5B, correlative cost of synthesis). There was a modest effect of feeding treatment (ANOVA, *F*_*1,32*_ = 5.29, *P* = 0.028, Table 1). However, there was no significant interaction between treatment and larval age (ANOVA, *F*_*1,32*_ = 2.16, *P* = 0.152, Table 1), indicating that the slopes of the regression between metabolic rate and protein synthesis rate for low- and high-fed larvae were similar. The treatment effect therefore is a result of a difference in the y-intercept of this relationship. As this energetic analysis uses only the slope of the relationship to estimate cost, a pooled regression for low- and high-fed larvae was used to determine the cost of protein synthesis. The common slope for both low- and high-fed is displayed as the solid line on Fig. 5B. This slope (which represents the amount of energy used per ng protein synthesized) results in a cost of synthesis of 5.34 ± 0.51 J (mg protein synthesized)^−1^ (± standard error of the slope, N = 33). For reference, the regressions for low- and high-fed are also shown (Fig. 5B, dashed lines), indicting similar slope values. The costs of synthesis for low- and high-fed separately were 4.29 and 5.58 J (mg protein synthesized)^−1^, respectively.

#### Percent Metabolism and Depositional Efficiency

Primary data relating to rates of protein growth (Fig. 2A), protein synthesis (Fig. 4A), and respiration (Fig. 2C) were used to model the protein depositional efficiency and the amount of metabolism used to support protein synthesis in low- and high-fed larvae. Protein depositional efficiency (PDE) (Fig. 6A) exhibited a trend that was similar to that observed in other physiological efficiencies (e.g., compare with Fig. 2D). Low-fed larvae had an initial efficiency that was 1.4-times higher than high-fed (83% and 60%, respectively). PDE decreased quickly in low-fed larvae while increasing modestly in high-fed larvae. At 24 DPF PDE was 1.6-times higher in high-fed than low-fed (66% and 41%, respectively).

**Figure 6.**
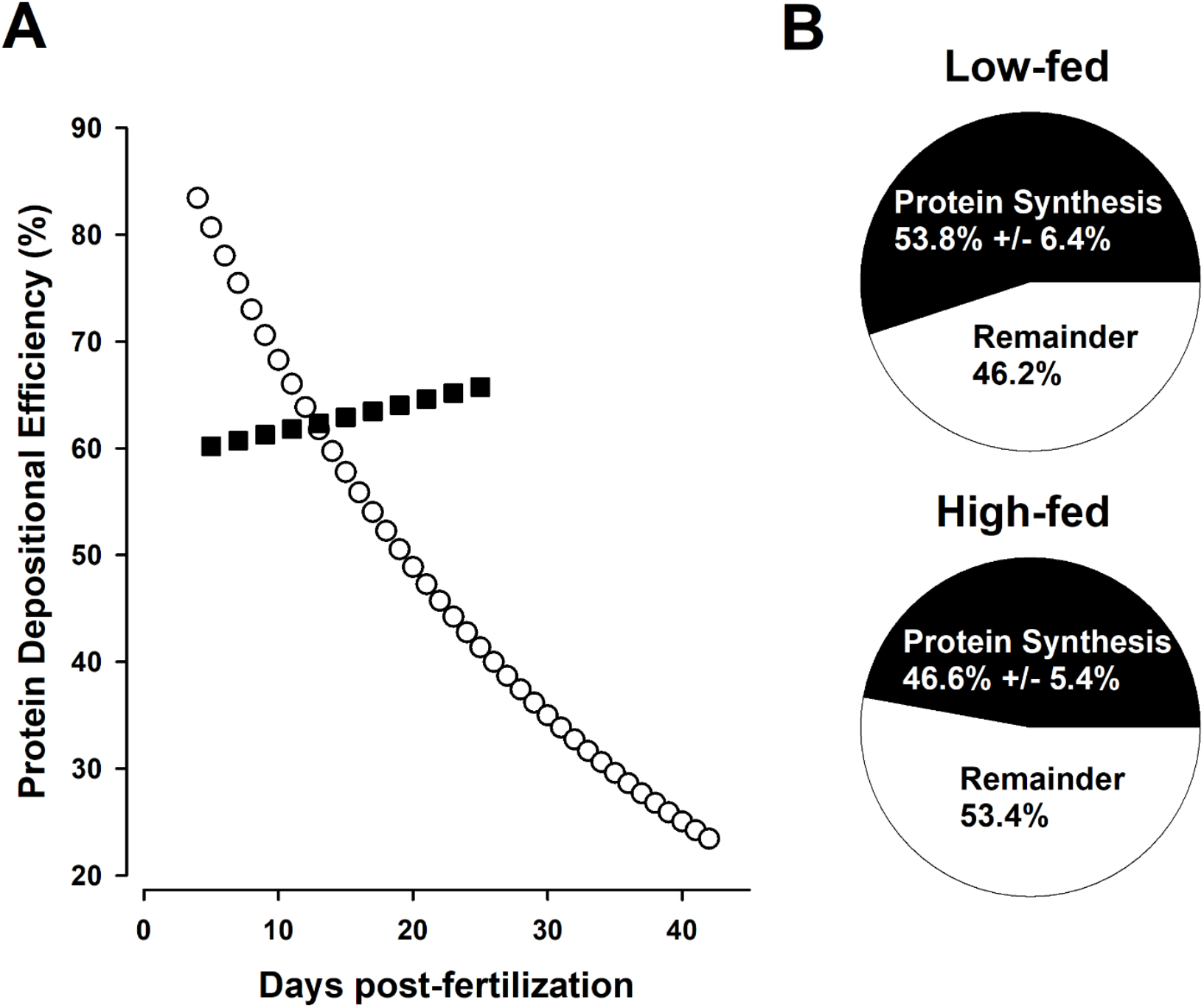
Protein depositional efficiency and percent metabolism of protein synthesis. **(A)** Modeled changes in Protein Depositional Efficiency (PDE). PDE is the percent of synthesized protein (taken from Fig. 4A) that is retained as larval biomass (taken from Fig. 2A). White circles = low-fed, black squares = high-fed. **(B)** Percent of metabolism used to fuel measured rates of protein synthesis. Protein synthesis rates (Fig. 4A) were multiplied by the grand average of the cost of protein synthesis (4.91 J mg^−1^ protein) to calculate the energy used to sustain protein synthesis rates. These values were then divided by the total metabolic rate (Fig. 2C) to return the percent of metabolic rate used to drive protein synthesis. Percent metabolism for protein synthesis was similar between low- and high-fed larvae (ANOVA, F_1,32_ = 0.59, *P* = 0.45).

To estimate the percent of metabolism used to support measured rates of protein synthesis, an average cost of protein synthesis was calculated at 4.91 J (mg protein synthesized)^−1^. This cost is the average of estimates from the inhibitor and correlative approaches (4.47 and 5.34 J (mg protein synthesized)^−1^, respectively). The average percent of metabolism supporting protein synthesis rates were similar between feeding treatments (ANOVA, *F*_*1,32*_ = 0.59, *P* = 0.45) at about 50% (Fig. 6B).

## Discussion

The goal of the current study was to define strategies relating to protein metabolism that are employed by organisms growing at different rates as a result of differing food environments. To do this, we have extended our analysis of physiological plasticity in larvae of the sand dollar, *D. excentricus*. Our previous work (Rendleman et al., 2018) examining energetic growth efficiencies, using the same conditions as in this study, implicated significant differences in the synthetic efficiency and/or the energetic cost of protein metabolism. The results of this study specifically test these ideas and show that there are both significant differences in protein metabolism as well as important similarities between larvae growing at different rates. These data provide empirical evidence that organisms can respond to different food environments by exploiting different macromolecular turnover strategies to achieve growth.

### Protein metabolism and differential growth during early development

The most important result of this study was that the large-scale differences in growth rate between low- and high-fed larvae were achieved by differences in mass-specific rates of protein degradation and not by differences in mass-specific rates of protein synthesis. While there were major differences in organismal rates of protein synthesis between low- and high-fed larvae, these differences were a result of high-fed larvae having a higher biomass than low-fed larvae. This is to be expected given the 10-fold difference in food availability. However, once synthesis rates were standardized to protein biomass (i.e., fractional rates of protein synthesis), rates of synthesis were similar between feeding treatments. Fractional rates of synthesis in this study of ~ 0.5-1 % hr^−1^ were similar to those for other echinoderm larvae experiencing positive growth (Pace and Manahan, 2006, 2007b; Ginsburg and Manahan, 2009).

The significant differences in protein depositional efficiency (PDE) between low- and high-fed larvae were an outcome of the treatment-specific differences in protein degradation. These differences in PDE provide at least a partial explanation for the large differences that were observed in protein growth efficiency (PGE) in both our original study of physiological plasticity (Rendleman et al., 2018) and the current study. With low-fed larvae being able to deposit more synthesized tissue as biomass (in early development), this results in high PGE and PDE values. Given that protein is the predominant biochemical substrate of these larvae (Rendleman et al., 2018), these differences in PDE can be viewed as the driving force for the differential growth efficiencies in these larvae when experiencing different food levels. A recent study employing similar techniques for measuring protein metabolism examined genotype-dependent differences in growth among different families of bivalve oyster larvae (Pan et al., 2018). Interestingly, the authors found growth differences to be related to differences in protein synthesis rates rather than changes in protein depositional efficiency. While comparison between the two studies is difficult due to our study manipulating environment and Pan et al. (2018) manipulating genotype to alter growth, as well as the difference in species studied, both studies demonstrate the important role that protein metabolism exerts on overall growth and development in feeding larval forms.

No significant differences were observed in the energetic cost of protein synthesis or its metabolic regulation in low- and high-fed larvae. As protein synthesis is typically the single-most expensive metabolic process (Hawkins, 1991; Houlihan, 1991), we used two techniques to arrive at an accurate energetic estimate of this critical cost. Both estimates returned similar energetic values with an average cost of 4.91 J (mg protein synthesized)^−1^. The first method relied on the eukaryotic translation inhibitor, anisomycin. This analysis allows for measuring costs in different feeding treatments at specific developmental times. A previous study has shown anisomycin to be both a specific and potent inhibitor of protein synthesis in larval echinoids (Pace and Manahan, 2007a). Based on the measured amount of inhibition and its lack of effect on alanine transport rates, our current analysis supports these previous findings. The second approach used a correlative method in which the average cost of synthesis over the entire range of larval development was estimated. We observed similar energetic costs of protein synthesis with respect to technique employed and with regards to feeding treatment and developmental stage. Furthermore, our energetic determinations of cost are similar to previous studies on larval echinoderms (Pace and Manahan, 2006, 2007a; Pan et al., 2015). Because we observed no significant differences in energetic cost, we could use this value to examine if the metabolic regulation of protein synthesis was feeding treatment specific. Here too, no significant differences were observed, and the average percent of metabolic energy used to support measured rates of synthesis was ~ 50%. This value of ~ 50% is in agreement with other studies of marine invertebrate larvae (Pace and Manahan, 2007a; Lee et al., 2016; Pan et al., 2018).

While rates of protein degradation were the major difference in growth efficiency between low- and high-fed larvae, the associated costs of degradation did not appear to be a significant factor in the plasticity response. The amount of metabolism required to support degradation rates increased for low-fed larvae over development, however, the estimated percent of metabolism was no more than 2%. These costs were estimated using the approach of Pan et al. (2018) relying on the cost of degradation measured by Peth et al. (2013). These estimates are for proteasomal degradation and likely have a significant amount of variance due to the folding patterns of degraded proteins, their level of ubiquitination, as well as differential costs associated with other degradation pathways. Even so, the total contribution appears to be relatively small, meaning the burden of high degradation rates is related to the high cost of resynthesizing those proteins that were degraded.

With both treatments having similar mass-specific rates of synthesis, the same energetic cost of synthesis, and utilizing a similar proportion of metabolism to support those rates, it seems likely that there are significant differences in the types of proteins that are being made. Given the 10-times higher food availability, high-fed larvae are likely making proportionally more structural proteins that can be deposited as biomass to support larval development as well as the creation of the juvenile rudiment. With significantly less food, low-fed larvae are likely making a proportionally higher number of metabolic enzymes serving regulatory purposes (i.e., house-keeping processes). These enzymes typically experience higher rates of turnover and would be a significant contributor to the relatively higher degradation rates. Importantly, these degradation rates increase over time and result with low-fed larvae having PDE values that are ~3-times less than high-fed by the end of development.

### Physiological mechanisms of developmental plasticity

Previous studies have indirectly implicated the role of protein metabolism in food-induced developmental plasticity in echinoid larvae (Heyland and Hodin, 2004; Carrier et al., 2015; Rendleman et al., 2018). For example, Heyland and Hodin (2004), demonstrated that exogenous thyroxine treatment can recapitulate the high-fed morphological phenotype (short arms, rapid rudiment development). Thyroxine is known to influence protein turnover rates (Sokoloff and Kaufman, 1961; Bates and Holder, 1988). In a transcriptomic study, Carrier et al. (2015) found evidence for the role of TOR (target of rapamycin) for regulating growth and development. TOR is known to be a master regulator of protein synthesis, with high levels causing an increase in ribosome biogenesis and subsequent up-regulation of translational processes (Raught et al., 2001). The direct measurements of protein metabolism in this study support these previous studies. The action of TOR may provide an explanation for why high-fed larvae are able to maintain equivalent levels of mass-specific protein synthesis despite their very rapid increase in protein biomass. TOR is also known to negatively regulate macroautophagy. For example during invertebrate larval development, TOR levels decrease when there is nutritional stress, increasing rates of macroautophagy (Scott et al., 2004; Di Bartolomeo et al., 2010; Romanelli et al., 2014). This provides an explanation for the higher overall degradation rates in low-fed as development proceeds, thereby allowing for the synthesis of high-turnover proteins. Adams et al. (2011) demonstrated that dopaminergic receptors respond to food environment and appear to cause the induction of the short-armed, high-fed larval phenotype. Due to the wide range of effects that dopamine can have on cellular physiology, it is difficult to know, at this time, what role it might play with regards to protein metabolism. Dopamine signaling is known to influence cellular plasticity by changing aspects of protein metabolism (Hasselgren et al., 1983; Smith et al., 2005; Pfeiffer and Huber, 2006). In *C. elegans* dopamine signaling results in higher levels of proteasomal degradation by increasing polyubiquitination of protein substrates (Joshi et al., 2016). This dopaminergic response, which is located in the epithelial cells of the worm, is activated when the worm encounters a bacterial lawn for feeding. Given the highly specific location of dopamine receptors at the tips of the pluteus larva’s post oral arms (Adams et al., 2011), this raises the possibility that the induction of the short-armed phenotype may be driven by increased degradation rates at the larval arms. This could then allow tissue growth to be focused towards post-larval structures such as the stomach and rudiment. These results are consistent with the increases seen in PDE as high-fed larvae develop and could be the consequence of increasing levels of TOR. Future analyses testing for such morphologically specific differences in protein homeostasis, as well as examining specific pathways of protein degradation (proteasomal vs autophagy) will be important for further understanding the role of both TOR and dopamine signaling mechanisms in the plasticity response.

Our analysis of protein metabolism also indicated that amino acid transport may play a critical role in the developmental plasticity response. Low-fed larvae had rates of alanine transport that were ~2-times higher than high-fed larvae. This result is of interest because it is likely related to the morphological plasticity response. The long-armed phenotype of low-fed larvae has been interpreted as a means to increase their feeding ability on algal cells (Hart and Strathmann, 1994). Meyer and Manahan (2009) found that amino acid transporters are primarily localized to the external epithelium of pluteus larvae, including the larval arms. The transport of dissolved organic material (DOM), such as alanine, has been shown to be an important contributor to acquiring energetic substrates for supporting larval metabolism (Manahan, 1990; Shilling and Manahan, 1994). Therefore, a long-armed phenotype may also serve the purpose of allowing greater access to the dissolved organic material (DOM) pool by way of increasing the epithelial surface area. It is also possible that low-fed larvae increase the epithelial density of DOM transporters. Such transporter mechanisms could also act as an environmental sensing mechanism by which plasticity responses are induced in prefeeding stages.

In total, the findings from other echinoid developmental plasticity studies in combination with the current results suggest an integrated response to food levels involving physiological and morphological responses which complement one another. Food conditions are sensed prior to the onset of feeding ability (Miner, 2007; Adams et al., 2011). If the larva is in a high food environment, then dopamine signaling will cause the induction of the short-armed phenotype (Adams et al., 2011). Part of this response may involve changes in protein turnover which have a negative influence on arm length and allow greater growth in post-larval structures to be supported. If food levels are low, then the default pathway of long larval arms is followed. The longer arms allow increased feeding ability (Hart and Strathmann, 1994) and DOM transport rates (this study). The increased algal feeding is supported by higher digestive efficiency (Rendleman et al., 2018 and this study). This increased digestive efficiency is further enhanced by increased protein depositional efficiency (this study), which is supported by the down-regulation of protein degradation. It is of interest that the default pathway elicits greater initial protein metabolic and growth efficiency. Understanding the molecular mechanisms that allow for high efficiency growth and comparing them to larval forms that do not have developmental plasticity would be of interest.

Relating these morphological and physiological responses to their adaptive value for the entirety of larval development is critical to understanding the actual ecophysiological value of developmental plasticity. It has been observed that the differential arm length response occurs early in larval development (Boidron-Metairon, 1988; Sewell et al., 2004; Miner, 2007). Physiologically, large differences in both digestive efficiency and growth efficiency (Rendleman et al., 2018 and this study) that benefit low-fed larvae are also seen only in early larval development. These results may be indicative of the cost-benefit relationship between nutrient acquisition and surface area exposure. Longer arms might also result in the need for greater surface area ion regulation. Previous research has demonstrated that the Na^+^-K^+^-ATPase pump can consume ~40% of larval metabolic energy (Leong and Manahan, 1997). Therefore, there may be a limit to the extent, both in total arm length and in length of developmental time, that long arms are adaptive. The long-armed phenotype may provide its greatest benefit early in the larval stage so as to allow the larva to find a more supportive nutritional environment without incurring large costs for surface area exposure. After early larval development, the survival strategy may alter to one of overall metabolic down-regulation with specific decreases in protein metabolism relating to growth in order to save energy and await better feeding conditions. In studies examining protein synthesis in unfed echinoid larvae (i.e., receiving no algal food), metabolic regulation of protein synthesis decreased to 16-21% of metabolism (Pace and Manahan, 2006; Pan et al., 2015), allowing for the maintenance of other essential processes such as Na^+^-K^+^-ATPase. These plastic physiological strategies likely allow for the long-term survival observed in nutritionally-stressed larvae (e.g., Moran and Manahan, 2004; Carrier et al., 2015).

### Echinoid larvae as a model system for developmental plasticity

The importance of, as well as the need to better understand, developmental plasticity has been expressed by evolutionary biologists, molecular biologists, ecologists, and physiologists (e.g., DeWitt et al., 1998; Miner et al., 2005; Evans and Hofmann, 2012; Uller et al., 2020). The developmental biology of sea urchins has long attracted researchers from all of these fields, resulting in a powerful set of integrative biological information. In conjunction with this, echinoid larvae have been the subject of many studies regarding developmental plasticity (recently reviewed in McAlister and Miner, 2018). These studies have shed important light on the molecular, morphological, and physiological manifestation of developmental plasticity. By virtue of these combined datasets, continued efforts using this system will provide a highly informative and integrated platform for understanding how organisms respond to changes in other important environmental conditions (e.g., temperature, pH, pollutants).

A critical point that has been brought up in previous analyses regarding developmental plasticity in echinoids is the need for experimental replication. Sewell et al. (2004) observed significant within-treatment variation associated with culture chambers. It has been recommended that future studies employ experimental designs by which to assess and partition this variation from the among-treatment effects (Sewell et al., 2004; McAlister and Miner, 2018). This current study, as well as our previous study on growth efficiency (Rendleman et al., 2018) employed 3 independent cultures (e.g., spawned at different times of the year using different parents). This was especially important given the integrative nature of our physiological measurements (e.g., combining both metabolic rate and protein synthesis rates) which can result in relatively large amounts of variance. We did observe significant within-treatment variation for both ingestion rates and respiration rates. However, this variation did not obscure the effect of food availability. In total, we now have six independent cultures, over the course of 4 years (initiated in different seasons), that show a consensus signal in the physiological strategies that are exploited to achieve positive growth in larvae experiencing different amounts of food. It is apparent that significant physiological plasticity, specifically relating to protein synthetic efficiency, accompanies the well-characterized morphological plasticity of these larvae. Future experiments will further explore this system to discover more regarding its mechanisms and adaptive value as well as identify its linkage to other important attributes of developmental plasticity such as TOR and dopamine signaling.

## Abbreviations

DPF: days post-fertilization
PO: postoral arm
AE: assimilation efficiency
GGE: gross growth efficiency
NGE: net growth efficiency
PGE: protein growth efficiency
PDE: protein depositional efficiency
MBL: midline body length
FAA: free amino acid pool
RGR: relative growth rate
BCA: bicinchoninic assay
BSA: bovine serum albumin
TCA: trichloroacetic acid
BOD: biological oxygen demand
TOR: target of rapamycin
DOM: dissolved organic material

## Acknowledgements

The authors thank Yvette Ralph for assistance in acquiring and maintaining adult sand dollars. We thank Kendra Ellis, Noah Grunfeld, Matan Grunfeld, and Annie Jean Rendleman for assistance with larval culturing and morphological and biochemical analyses. Dr. Bengt Allen assisted with statistical analyses. Dr. Bruno Pernet provided critical feedback on an earlier draft of this paper. This work was supported by the National Institute of General Medical Sciences of the National Institutes of Health under Award Numbers UL1GM118979, TL4GM118980, and RL5GM118978. The content is solely the responsibility of the authors and does not necessarily represent the official views of the National Institutes of Health. Funding was also provided by CSU-COAST (Council on Ocean Affairs, Science and Technology) and from the Los Angeles Rod and Reel Scholarship fund (awards to A. Ellison).

